# Surfaces: A software to quantify and visualize interactions within and between proteins and ligands

**DOI:** 10.1101/2023.04.26.538470

**Authors:** Natália Teruel, Vinicius Magalhães Borges, Rafael Najmanovich

**Affiliations:** Department of Pharmacology and Physiology, Faculty of Medicine, Université de Montréal, Montreal, Canada; Department of Biomedical Sciences, Joan C. Edwards School of Medicine, Marshall University, Huntington, WV, USA

## Abstract

Computational methods for the quantification and visualization of the relative contribution of molecular interactions to the stability of biomolecular structures and complexes are fundamental to understand, modulate and engineer biological processes. Here we present Surfaces, an easy to use, fast and customizable software for quantification and visualization of molecular interactions based on the calculation of surface areas in contact. Surfaces calculations shows equivalent levels of correlations with experimental data as computationally expensive methods based on molecular dynamics. All scripts are available at https://github.com/nataliateruel/Surfaces

Documentation is available at https://surfaces-tutorial.readthedocs.io/en/latest/index.html

## 1 Introduction

Molecular interactions determine all aspects of biological processes. Understanding the relative contributions of individual molecular interactions can guide the design of effective drugs to modulate biological processes as well as help understand the effect of mutations in natural processes or guide their introduction in protein engineering.

Protein engineering is an important tool in synthetic biology that can produce proteins with incredible therapeutic and industrial potential (Tobin *et al*., 2015; Chica, 2015; Bojar and Fussenegger, 2020). In recent years, the field has made giant strides forward, but it still has the potential for exponential growth – as seen for many fields that benefit from high throughput technologies and powerful new computational tools. However, a random search for all possible sequence configurations and respective structures and functions might reveal an enormous searchable space to explore. This is, even today, one of the biggest challenges of protein engineering, and one that makes the rational design extremely necessary. Techniques that provide insights into the partial contribution of each atom or residue to protein interactions are critical for understanding the intricacies of the interaction interface. By identifying the specific residues and atoms that play a crucial role in the interaction, these techniques can aid in the development of rational protein engineering strategies. Unfortunately, these techniques, mostly based on *ab initio* approaches, are often computationally expensive, limiting the extent to which they can be used to explore this expansive searchable space, like through exhaustive mutant modelling evaluations or large datasets.

With the decreasing cost of computational power, it is possible to simulate biological systems considering more realistically the underlying physical processes at the core of molecular interactions. Molecular dynamics (MD) simulations are at the forefront of such efforts, but such methods are computationally expensive and often also difficult to implement and thus remain impractical for high-throughput applications or broad adoption. Among these methods, we can highlight free-energy perturbation (FEP) methods that aim at predicting the DDG of binding for different mutations relative to wild-type or the binding DG for biomolecular complexes (Lavigne *et al*., 2000; Sergeeva *et al*., 2022; Zhu *et al*., 2022; McCarrick and Kollman, 1999) and gRINN (Serçinoglu and Ozbek, 2018), a tool based on MD simulations to breakdown the per-residue energetic contribution of molecular interactions.

To address this challenge, simplified techniques using atomic surface areas in contact to estimate binding energy can be employed. This approach has previously been introduced in methods such as LPC (<u>L</u>igand-<u>P</u>rotein <u>C</u>ontacts) and CSU (<u>C</u>ontacts of <u>S</u>tructural <u>U</u>nits) (Sobolev *et al*., 1999) as well as STC (<u>S</u>tructure-based <u>T</u>hermodynamic <u>C</u>alculations) (Lavigne *et al*., 2000). While these techniques are currently limited by server availability, as well as pre-defined atom type definitions and energetic pairwise matrices, they provide a faster path to evaluating energetic decomposition and served as the basis upon which many other methods that use atomic surfaces areas in contact were built (Frappier *et al*., 2017; Gaudreault and Najmanovich, 2015; Ribeiro *et al*., 2019; Olechnovič and Venclovas, 2021). Therefore, it is worth revisiting and improving these techniques by employing new libraries and visualization tools, making them customizable and user-friendly to broaden their general applicability. By doing so, we can enhance our understanding of the per-residue contribution to protein interactions and facilitate more efficient and effective protein engineering.

In this application note, we present Surfaces, a fast method that utilizes atomic surface areas in contact, user-defined atom-type definitions, and pairwise pseudo-energetic matrices as a proxy for highlighting favorable and unfavorable interactions within and between proteins, as well as between proteins and other biomolecules.

## 2 Methods

Surfaces quantifies atomic interactions using two measures. The first measure is the area in contact between atoms as described and calculated by Vcontacts using a Voronoi procedure (McConkey *et al*., 2002). This method restricts the evaluation of interactions to atoms within close proximity. The second measure is a pairwise pseudo-energetic matrix that assigns an interaction value based on the atom types (Fig 1A).

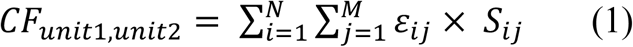

**Fig. 1.**
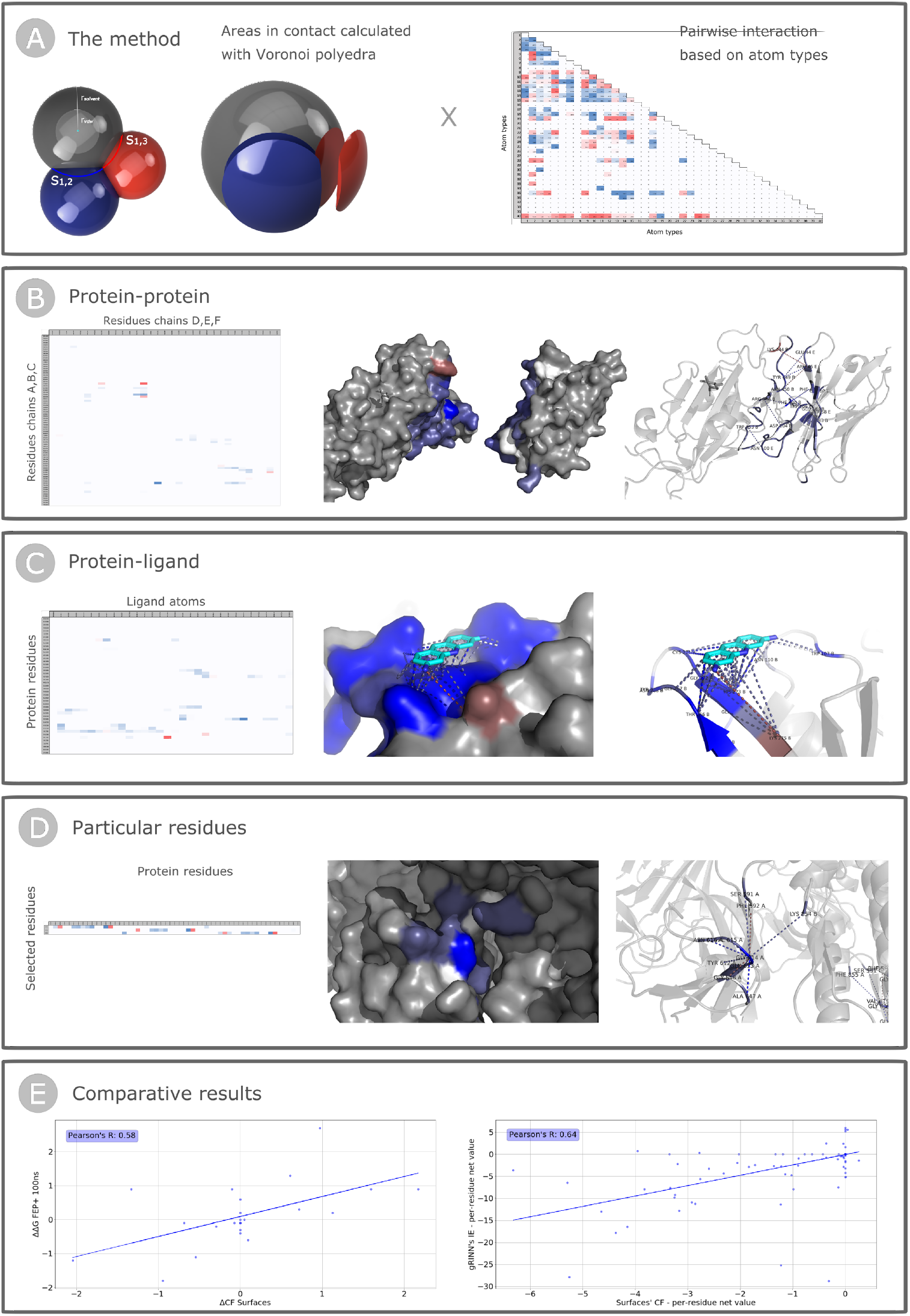
Schematic representation of the method and applications of Surfaces scripts. **(A)** The method: Representation of the calculated areas in contact between atoms 1 (gray), 2 (blue) and 3 (red), considering the expanded radii - that accounts for the van der Waals radii (*r*_*vdw*_) and the radius of a water molecule (*r*_*solvent*_) – and leading to the definition of the surfaces in contact *S*_1,2_ and *S*_1,3_, represented as exploded spherical caps, and the default matrix of pairwise interactions based on 40 SYBYL atom types. **(B)** Protein-protein: Example of the *protein-protein* application using the PDB structure 7VQ0 (Maeda *et al*., 2022). Representation of the .CSV output and of the visual output colored from red (unfavorable) to blue (favorable) showing the net value of interactions mapped into the surface of the chains, as well as the most relevant interactions highlighted with dash lines. **(C)** Protein-ligand: Example of the *protein-ligand* application using the PDB structure 7NT4 (Napolitano *et al*., 2022). Representation of the .CSV output and of the visual output colored from red (unfavorable) to blue (favorable) showing the net value of interactions mapped into the surface of the protein, as well as the interactions with the atoms of the ligand highlighted with dash lines. **(D)** Particular residues: Example of the *particular residue*s application using the PDB structure 7EAZ (Yang *et al*., 2021) and selecting the residues GLY614. Representation of the .CSV output and of the visual output colored from red (unfavorable) to blue (favorable) showing the net value of interactions mapped into the surface of the protein, as well as the interactions involving the residues of interest highlighted with dash lines. **(E)** Comparative results of Surfaces with the calculations performed using FEP + 100 ns MD simulations for the interaction of 23 mutants of the Spike protein and the receptor ACE2 (Sergeeva *et al*., 2022), and with the pairwise Interaction Energies calculated with gRINN (Serçinoglu and Ozbek, 2018) for a MD simulation of the Spike protein Receptor-Binding Domain of the Delta variant and the receptor ACE2 (Cheng *et al*., 2022).

The complementarity function (CF) that accounts for these two measures is described by equation 1, in which *N* is the total of atoms of unit 1, *M* is the total of atoms of unit 2, *ε*_*ij*_ is the energy for the interaction between atoms i and j, and *S*_*ij*_ is the surface area in contact between atoms *i* and *j*. Units can correspond to residues, ligands or ligand atoms.

The CF calculation is based on the scoring function utilized by FlexAID (Gaudreault and Najmanovich, 2015), a ligand docking tool that also provides the SYBYL definition of atom types and the respective matrix of pairwise interaction energies used as the default in Surfaces scripts, itself based on the LIGIN docking software (Sobolev *et al*., 1996). These scripts are designed to be customizable, enabling the incorporation of modified residues or ligands, as well as the use of alternative atom type classifications and energetic matrices.

To visually represent the results of interaction analysis, Surfaces offers functions created using the PyMOL API (DeLano). These functions are also customizable and allow for the visualization of the overall contribution of residues to interaction surfaces, as well as specific interactions between structural units, as PyMOL sessions.

## 3 Results and discussion

Surfaces was built as a set of python scripts designed to evaluate various types of interactions in proteins. The scripts are user-friendly, fast, and can be easily customized and automated, offering a scriptable way of generating data and visual representations for the interactions of interest.

### 3.1 Applications of Surfaces

The specific types of interactions that can be analysed by Surfaces are: Protein-protein interactions, protein-ligand interactions and residue interactions. In all cases, the input is a .PDB formatted file while the output of the quantitative analysis save as a .CSV file. The output for visualisation is saved as a .PSE PyMOL session file. The visual output shows the surfaces of the relevant units of interest colored according to the net value of interactions and the specific pairwise interactions to the net total are represented as colored dash lines with a customizable color scale. In all cases the input .PDB structure needs to be cleaned of any non-defined atoms in a pre-processing step, which can be done using scripts that are provided with the Surfaces software. The pre-processing is not done automatically to offer the use the possibility to customize the analysis. For example, if the user wishes to ignore specific hetero-atoms, these need to be removed but otherwise, defined.

#### Protein-protein interfaces

For this application, two groups of protein chains are required as input, Surfaces calculates the interactions between residues of the first group and residues of the second group (Fig 1B). This can also be used for protein-ligand evaluations by assigning a chain ID for the ligand’s atoms, if the user intends to evaluate the complete interaction of the ligand with each residue without a per-atom decomposition (see next paragraph).

#### Protein-ligand interactions

This application requires the PDB assigned 3-letter code of the ligand of interest as input and calculates the interaction between residues of the structure and each atom of every instance of the ligand within the structure (Fig 1C). Before running the application, two pre-processing steps are required: first, the assignment of atom types to the atoms of the ligand into the .DEF file, and second, cleaning the input .PDB structure of any non-defined atoms. Surfaces offers a general script for atom type assignment mapping element types to the default SYBYL 40 atom types.

#### Residue interactions

This application requires a list of residues of interest as input and calculates all interactions (inter- or intra-chain) involving those residues (Fig 1D).

### 3.2 Validation of Surfaces

The most common methods for analyzing protein-protein interactions and the per-residue energetic decomposition of the contributions of such interactions to the overall free energy are based on MD simulations (Homeyer and Gohlke, 2012; Kollman *et al*., 2000; Serçinoglu and Ozbek, 2018). MD simulations can provide detailed information on the structural changes and energetics associated with protein-protein interactions, including the binding free energy and per-residue energetic contributions. However, MD simulations are computationally intensive (Ciccotti *et al*., 2022; Bopp *et al*., 2008) making them less suitable for large-scale studies including protein engineering. Furthermore, MD simulations require expert knowledge to set up, which creates an additional obstacle for broad utilization.

The examples presented in Figure 1 to illustrate the utilization of Surfaces to the three applications of Surfaces were taken from different datasets. The *protein-protein* interface analyzed in Fig. 1A was taken from a dataset of 734 structures of one or more chains of the SARS-CoV-2 Spike protein in complex with one or more chains of antibodies (Gowthaman *et al*., 2020). Such an analysis requires an average of 11.38 ± 9.62 CPU-seconds per structure including the time required for the preprocessing step. The analysis of *protein-ligand* interactions is derived from a dataset of 709 non-redundant structures and 669 non-redundant ligands, totaling 831 experimentally solved protein-ligand complexes of SARS-CoV-2 proteins and small molecules (Harrison *et al*., 2021). This analysis requires an average of 3.97 ± 2.32 CPU-seconds per structure (including pre-processing).

To demonstrate the reliability of Surfaces, we compare its results with a highly curated dataset of 23 SARS-CoV-2 Spike Receptor Binding Domain (RBD)/ACE2 binding DDG values measured by Surface Plasmon Resonance (Sergeeva *et al*., 2022) and a compare these to large number of computational methods. Specifically: FEP + 100ns MD simulations From Sergeeva *et al*. (Sergeeva *et al*., 2022), methods based on machine-learning: Mutabind2 (Zhang *et al*., 2020), mCSM-PPI (Rodrigues *et al*., 2019), SAAMBE-3D (Pahari *et al*., 2020); based on statistical potentials: BeAtMusic (Dehouck *et al*., 2013); and force field related scoring functions: FoldX (Schymkowitz *et al*., 2005), Rosetta flex ddG (Barlow *et al*., 2018). Surfaces shows a Pearson’s correlation coefficient (PCC) of 0.556 with the experimental data, while FEP+100 ns MD obtains a PCC of 0.598. All other methods obtain considerably lower PCC values (Supplementary Table S1 and Supplementary Figure S1). Surfaces and FEP+100 ns obtain DDG root mean square errors (RMSE) of 0.747 and 0.754 kcal/mol respectively. The DDG RMSE for the other methods vary from 0.475 for SAAMBE-3D (PCC of 0.259) and 1.741 for FoldX (PCC of -0.079). The PCC between FEP+100 ns and Surfaces is 0.58 (Figure 1E).

A widely used method for residue-based energy decomposition is gRINN (Serçinoglu and Ozbek, 2018), which uses MD trajectories to calculate binding energy means and distributions. Using trajectories of Delta SARS-CoV-2 Spike in complex with the receptor ACE2 (Cheng *et al*., 2022), available at the COVID-19 Molecular Structure and Therapeutics Hub (MolSSI), we used gRINN to calculate interface interactions and compare them to Surfaces results. The Pearson’s correlation coefficient obtained comparing Surfaces and gRINN is 0.64 (Figure 1E and Supplementary Figure S2).

In order to consider structural variations, captured when using MD-based approaches, Surfaces can be applied to protein ensembles, generated with tools such as the NRGTEN package (Mailhot and Najmanovich, 2021). This allows for the evaluation of transient interactions, as well as the calculation of the energy distribution.

## 4 Conclusions

Surfaces provides a simple, fast and easy to use method to analyze and visualize biomolecular interactions that is as effective computationally expensive and cumbersome to implement methods such as gRINN and FEP. The use of variations in solvent accessible surface areas or those in contact to estimate free-energy contributions has been stablished previously (with LPC/CSU, STC) but somehow fell out of use over the past 20 years within the broader community. However, this work shows that such calculations can provide relevant insight of the same level than that obtained using more computationally expensive methods based on molecular dynamics or even more than currently available methods based on machine learning or other mean-field based statistical approaches.

The scriptable and customizable nature of Surfaces makes it a valuable tool for researchers seeking to analyze large structural datasets, such as in virtual screening or protein engineering, thus allowing a broader exploration of search space leading to a narrowing down of alternatives prior to more computationally expensive computational analyses or experiments. Lastly, the ease of use of Surfaces makes this type of analysis accessible to a broader audience.

## Acknowledgements

We would like to thank Nicolas DesCôteaux for the thorough testing of the software and identification of initial bugs. This work is dedicated to Vladimir Sobolev (Z”L, 1947-2023), creator of LIGIN and LPC/CSU and a pioneer in the utilization of surface areas in contact as a fundamental component of the analysis of biomolecular interactions.

## References

Barlow, K.A. et al. (2018) Flex ddG: Rosetta Ensemble-Based Estimation of Changes in Protein– Protein Binding Affinity upon Mutation. J Phys Chem B, 122, 5389–5399.

Bojar, D. and Fussenegger, M. (2020) The Role of Protein Engineering in Biomedical Applications of Mammalian Synthetic Biology. Small, 16, 1903093.

Bopp, P.A. et al. (2008) SCOPE AND LIMITS OF MOLECULAR SIMULATIONS. Chem Eng Commun, 195, 1437–1456.

Cheng, M.H. et al. (2022) Impact of new variants on SARS-CoV-2 infectivity and neutralization: A molecular assessment of the alterations in the spike-host protein interactions. Iscience, 25, 103939.

Chica, R.A. (2015) Protein engineering in the 21st century. Protein Sci, 24, 431–433.

Ciccotti, G. et al. (2022) Molecular simulations: past, present, and future (a Topical Issue in EPJB). Eur. Phys. J. B, 95, 3.

Dehouck, Y. et al. (2013) BeAtMuSiC: prediction of changes in protein–protein binding affinity on mutations. Nucleic Acids Res, 41, W333–W339.

DeLano, W. The PyMOL Molecular Graphics System. Schrödinger, LLC. (www.pymol.org).

Frappier, V. et al. (2017) Applications of Normal Mode Analysis Methods in Computational Protein Design. Methods in molecular biology (Clifton, NJ), 1529, 203–214.

Gaudreault, F. and Najmanovich, R.J. (2015) FlexAID: revisiting docking on non-native-complex structures. Journal of chemical information and modeling, 55, 1323–1336.

Gowthaman, R. et al. (2020) CoV3D: a database of high resolution coronavirus protein structures. Nucleic Acids Res, 49, gkaa731..

Harrison, P.W. et al. (2021) The COVID-19 Data Portal: accelerating SARS-CoV-2 and COVID-19 research through rapid open access data sharing. Nucleic Acids Res, 49, gkab417..

Homeyer, N. and Gohlke, H. (2012) Free Energy Calculations by the Molecular Mechanics Poisson−Boltzmann Surface Area Method. Mol Inform, 31, 114–122.

Kollman, P.A. et al. (2000) Calculating Structures and Free Energies of Complex Molecules: Combining Molecular Mechanics and Continuum Models. Accounts Chem Res, 33, 889–897.

Lavigne, P. et al. (2000) Structure-based thermodynamic analysis of the dissociation of protein phosphatase-1 catalytic subunit and microcystin-LR docked complexes. Protein Sci, 9, 252–264.

Maeda, R. et al. (2022) A panel of nanobodies recognizing conserved hidden clefts of all SARS-CoV-2 spike variants including Omicron. Commun Biology, 5, 669.

Mailhot, O. and Najmanovich, R. (2021) The NRGTEN Python package: an extensible toolkit for coarse-grained normal mode analysis of proteins, nucleic acids, small molecules and their complexes. Bioinformatics, 37, 3369–3371.

McCarrick, M. and Kollman, P. (1999) Predicting relative binding affinities of non-peptide HIV protease inhibitors with free energy perturbation calculations. Journal of Computer-Aided Molecular Design, 13, 109–121.

McConkey, B.J. et al. (2002) Quantification of protein surfaces, volumes and atom-atom contacts using a constrained Voronoi procedure. Bioinformatics, 18, 1365–1373.

MolSSI COVID-19 Molecular Structure and Therapeutics Hub. Napolitano, V. et al. (2022) Acriflavine, a clinically approved drug, inhibits SARS-CoV-2 and other betacoronaviruses. Cell Chem Biol, 29, 774-784.e8.

Olechnovič, K. and Venclovas, Č. (2021) VoroContacts: a tool for the analysis of interatomic contacts in macromolecular structures. Bioinformatics, 37, 4873–4875.

Pahari, S. et al. (2020) SAAMBE-3D: Predicting Effect of Mutations on Protein–Protein Interactions. Int J Mol Sci, 21, 2563.

Ribeiro, J. et al. (2019) Calculation of accurate interatomic contact surface areas for the quantitative analysis of non-bonded molecular interactions. Bioinformatics, 35, btz062.

Rodrigues, C.H.M. et al. (2019) mCSM-PPI2: predicting the effects of mutations on protein– protein interactions. Nucleic Acids Res, 47, W338–W344.

Schymkowitz, J. et al. (2005) The FoldX web server: an online force field. Nucleic acids research, 33, W382–8.

Serçinoglu, O. and Ozbek, P. (2018) gRINN: a tool for calculation of residue interaction energies and protein energy network analysis of molecular dynamics simulations. Nucleic acids research, 46, W554–W562.

Sergeeva, A.P. et al. (2022) Free energy perturbation calculations of mutation effects on SARS-CoV-2 RBD::ACE2 binding affinity. Biorxiv, 2022.08.01.502301.

Sobolev, V. et al. (1999) Automated analysis of interatomic contacts in proteins. Bioinformatics, 15, 327–332.

Sobolev, V. et al. (1996) Molecular docking using surface complementarity. Proteins, 25, 120–129.

Tobin, P. et al. (2015) Protein Engineering: A New Frontier for Biological Therapeutics. Curr Drug Metab, 15, 743–756.

Yang, T.-J. et al. (2021) D614G mutation in the SARS-CoV-2 spike protein enhances viral fitness by desensitizing it to temperature-dependent denaturation. J Biol Chem, 297, 101238.

Zhang, N. et al. (2020) MutaBind2: Predicting the Impacts of Single and Multiple Mutations on Protein-Protein Interactions. Iscience, 23, 100939.

Zhu, F. et al. (2022) Large-scale application of free energy perturbation calculations for antibody design. Sci Rep-uk, 12, 12489.

