## Supplementary Figures and Tables for "Surfaces: A software to quantify and visualize interactions within and between proteins and ligands"

*To whom correspondence should be addressed.

**Binding evaluation through ΔΔG estimation**

Many different methods have been developed, based on different strategies, to quantify the binding affinity of protein-protein interfaces. Sergeeva *et al.* (Sergeeva *et al.*, 2022) compares their results using free-energy pertubations (FEP) against a broad range of different methods, specifically methods based on machine-learning: Mutabind2 (Zhang *et al.*, 2020), mCSM-PPI (Rodrigues *et al.*, 2019), SAAMBE-3D (Pahari *et al.*, 2020); based on statistical potentials: BeAtMusic (Dehouck *et al.*, 2013); and force field related scoring functions: FoldX (Schymkowitz *et al.*, 2005), and Rosetta flex ddG (Barlow *et al.*, 2018). The authors compare the performance of the different methods on a set of SARS-CoV-2 Spike mutations for which they have experimentally determined binding affinity to the receptor ACE2 using surface plasmon resonance (Pattnaik, 2005). Their data allows us to compare the performance of Surfaces to that of the various methods against experimental results.

To do so, we performed all the calculations on mutants modeled to the same crystal structure of ACE2/RBD (PDB 6M0J) (Lan *et al.*, 2020), following the same methodology used by Sergeeva *et al.* (Sergeeva *et al.*, 2022). Surface's CF scoring for the full binding interface was calculated by summing all individual interaction contributions from the per-residue outputs. The CF calculations Surfaces generates are based on a pseudo-physical model that does not follow the same kcal/mol scale used for the other methods. In order to do the RMSE calculation, as well as to follow particular numerical thresholds used for the analysis of the data, we also produced Surfaces results subjected to a linear regression to the experimental values.

The statistical metrics originally used to compare the results using the different methods were the Pearson's correlation coefficient (PCC), root mean square error (RMSE) and Pearson's phi for stabilizing mutations, performing a binary classification of the data, considering as stabilizing the mutations with $\Delta\Delta G \leq -0.4$ (PCC $\Phi$ (stabilizing $\leq$ -0.4)), an arbitrary threshold defined by Sergeeva *et al.* (Sergeeva *et al.*, 2022). We utilized the same statistical metrics to Surface results. The results are shown in Table S1. In Figure S1A we plot the experimental vs. calculated ΔΔG values for all the methods presented by Sergeeva *et al.* and in Figure S1B for Surfaces. According to all metrics, Surfaces' performance is equivalent to that of FEP, with a superior PCC to all other methods at a fraction of the computational cost.

It is worth noting that Surfaces provides a simplified estimation of changes in enthalpy differences (ΔΔH), not of the free energy. In the past we have utilized our ENCoM normal mode analysis method to estimate ΔΔG of mutations based solely on vibrational entropy differences (Frappier and R. J. Najmanovich, 2014; Frappier and R. Najmanovich, 2014). Such vibrational entropy differences, as calculated for a large number of Spike mutants (Teruel *et al.*, 2021), can be performed in a straightforward manner with NRGTEN package (Mailhot and Najmanovich, 2021) and combined with Surfaces ΔΔH predictions.

**Table S1. Comparative measures of binding affinity of mutants of the SARS-CoV-2 RBD and the receptor ACE2.** From Sergeeva *et al.*, 2022 we have the experimental values (ΔΔG experiment SPR), as well as the predictions using different binding affinity calculation methods (left table) and the analysis of Pearson's correlation coefficient (PCC), root mean square error (RMSE) and Pearson's phi for stabilizing mutations (PCC Φ (stabilizing ≤ -0.4)) (lines in blue). Added to this table, ΔCF Surface results calculated for the same mutants, the linear regression of these values to the experimental reference (right table) and the statistical analysis for Surfaces results (yellow lines). Green values correspond to stabilizing mutations with ΔΔG ≤ -0.4. Orange values correspond to destabilizing mutations with ΔΔG ≥ 0.4.

| RBD mutation | ΔΔG experiment SPR | ΔΔG Mutabind2 | ΔΔG mCSM-PPI | ΔΔG SAAMBE-3D | ΔΔG BeAtMusic | ΔΔG FoldX | ΔΔG Rosetta flex ddG | ΔΔG FEP+ 100ns | ΔCF Surfaces | ΔCF Surfaces (linear regression) |
|---|---|---|---|---|---|---|---|---|---|---|
| N501Y | -0.8 | 0.7 | 0.5 | 0 | 0.1 | 6 | 0.4 | -1.2 | -2837.6442 | -2.0 |
| Y453F | -0.7 | -0.2 | 0.1 | 0.1 | 0.3 | -0.4 | -0.2 | -0.6 | 128.30585 | 0.1 |
| S477N | -0.5 | -0.1 | -0.1 | 0.2 | 0.1 | 0 | 0.1 | -0.1 | -951.0603 | -0.7 |
| N501T | -0.5 | -0.6 | -0.9 | -0.1 | 0.4 | -0.9 | -0.4 | -1.8 | -1312.2174 | -0.9 |
| N439K | -0.1 | -0.1 | 0.5 | 0.6 | 0.3 | 0 | 0 | 0.6 | 0 | 0.0 |
| N440K | 0 | 0.1 | -0.1 | 0.5 | 0 | -0.1 | 0 | -0.4 | 0 | 0.0 |
| F490S | 0 | 0.6 | 0.2 | 0.8 | 0.7 | 0 | 0 | -0.1 | -104.3611 | -0.1 |
| L452M | 0 | 0 | 0.5 | -0.3 | 0.2 | 0 | 0 | 0.2 | 0 | 0.0 |
| L452R | 0 | -0.8 | 0 | 0.1 | 0.2 | -0.3 | 0 | -0.3 | 0 | 0.0 |
| E484Q | 0.1 | 0.1 | 0.3 | 0.5 | 0.2 | -0.1 | 0 | 0.3 | 993.9054 | 0.7 |
| T478K | 0.1 | 0.2 | 0 | 0.1 | 0.2 | 0 | 0.1 | 0 | 0 | 0.0 |
| N481K | 0.1 | 0 | 0 | 0.6 | 0.2 | 0 | 0 | -0.1 | 0 | 0.0 |
| E484K | 0.1 | 0.2 | 0.3 | 0.3 | 0.1 | -0.2 | -0.1 | 0.2 | 1564.06845 | 1.1 |
| Q498R | 0.2 | 0.4 | 1.4 | 1.1 | 0.8 | -0.5 | -0.6 | 2.7 | 1345.49484 | 1.0 |
| S477I | 0.2 | 0.1 | 0 | 0 | 0.2 | 0.1 | 0 | 0 | 68.5691 | 0.0 |
| G446V | 0.2 | -0.7 | 0.1 | -0.3 | 1.3 | 0.1 | 0 | 0.9 | -145.36265 | -0.1 |
| T478R | 0.2 | 0.1 | 0 | 0.3 | 0.2 | 0 | 0 | -0.1 | 0 | 0.0 |
| S477R | 0.3 | 0.2 | -0.1 | 0 | 0.1 | 0 | -0.2 | -0.2 | -413.462 | -0.3 |
| A475V | 0.3 | 0.9 | 0 | 0.1 | 0.1 | 1 | 0.2 | -1.1 | -760.3262 | -0.5 |
| L455F | 0.4 | 1.5 | -0.7 | 0.2 | -0.1 | 5 | -0.6 | 0.9 | -1850.59475 | -1.3 |
| K417T | 0.4 | 0.2 | 0.4 | 0.5 | 0.5 | 0.8 | 0.7 | 0.9 | 3011.74485 | 2.2 |
| F486L | 0.6 | 0.3 | 0.9 | 0.5 | 0.7 | 1.1 | 1.8 | 1.3 | 842.9372 | 0.6 |
| K417N | 0.6 | 0.6 | 0.5 | 0.4 | 0.5 | 0.8 | 0.9 | 0.9 | 2210.627225 | 1.6 |
| PCC | | 0.322 | 0.218 | 0.259 | 0.219 | -0.079 | 0.344 | 0.598 | | 0.556 |
| RMSE | | 0.523 | 0.532 | 0.475 | 0.497 | 1.741 | 0.501 | 0.754 | | 0.747 |
| PCC Φ (stabilizing ≤ -0.4) | | 0.163 | 0.266 | - | - | 0.504 | 0.163 | 0.592 | | 0.593 |

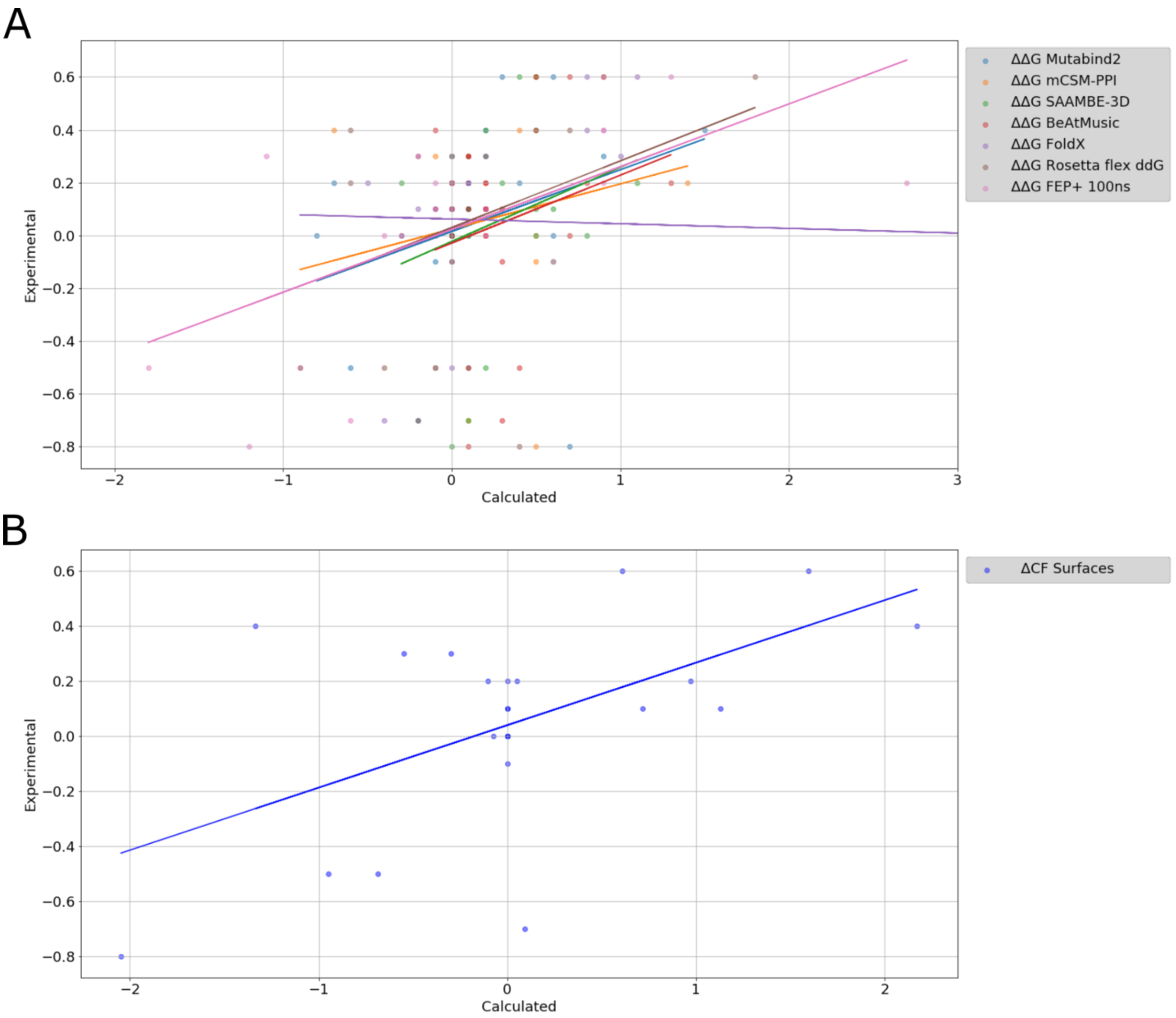


**Fig S1. Graphic representation of the calculated binding affinities for each of the mutants against the experimental values, as well as their tendency lines. (A)** Values of ΔΔG predictions using different binding affinity calculation methods compared to experimental data (Sergeeva *et al.*, 2022). **(B)** Linear regression of Surfaces' ΔCF prediction values for binding affinity compared to experimental data.

According to the different metrics used to compare predictive performance, FEP calculations based on a 100ns Molecular Dynamics (MD) trajectories are shown to represent a good alternative (Table S1). As noted by the authors (Sergeeva *et al.*, 2022), being MD-based, FEP calculations require a complex computational infrastructure and are time consuming. Surfaces, presenting a close Pearson's correlation, similar error compared to the experimental values, as well as a comparable classifying performance, measured by the Pearson's phi according to the determined threshold used for classifying a mutation as "stabilizing" (Table S1, Fig S1), required a very small fraction of the computational power to evaluate all mutants (185 CPU-seconds). The results from Table S1 and Fig S1 were generated with the goal to understand binding affinities of mutations and variants already selected during viral evolution. However, some of these same methods, with significant computational cost, would not be suitable for exploratory objectives while Surfaces can be used in high-throughput computational mutational scans.

While the measure of full binding of mutants is often used to evaluate the impact of mutations on protein interactions, it does not always reflect the changes in interactions of the mutated residue alone, and may be associated with a disruption of adjacent regions in the interface. This point is also discussed when observing the non-cumulative effects seen for single mutants on full interfaces – the sum of the effects of individual mutations may not equal the effect of multiple mutations combined – as pointed out in the first evaluations of the Omicron Spike (Cameroni *et al.*, 2022; Dejnirattisai *et al.*, 2022).

Due to these limitations, for exploratory objectives and rational design, per-residue energetic decomposition methods offer very important information for purposes of protein engineering.

**Per-residue decomposition**

The most common methods for analyzing protein-protein interactions and per-residue energetic decomposition are based on molecular dynamics (MD) simulations (Homeyer and Gohlke, 2012; Kollman *et al.*, 2000; Serçinoglu and Ozbek, 2018). MD simulations can provide detailed information on the structural changes and energetics associated with protein-protein interactions, including the binding free energy and per-residue energetic contributions. However, MD simulations are computationally intensive and therefore can take a long time to complete (Ciccotti *et al.*, 2022; Bopp *et al.*, 2008), making them less suitable for large-scale studies such as in computational protein design.

A widely used method for residue-based energy decomposition is gRINN (Serçinoglu and Ozbek, 2018), which uses MD trajectories to calculate binding energy means and distributions. From trajectories of the SARS-CoV-2 Spike Delta variant in complex with the receptor ACE2 (Cheng *et al.*, 2022), available at the COVID-19 Molecular Structure and Therapeutics Hub (MolSSI), we calculated interface interactions using gRINN and compare with Surfaces. We generated the results for gRINN considering only residues that were at a maximum distance of 6.8Å from the interface, a filtering distance of 20Å between residues and interactions present in 100% of the trajectory frames – these parameters reduced gRINN analysis to interactions in closer proximity, inter-chain interacting residues within the interface and filtered off transient interactions, to more closely match the interaction that are analyzed with Surfaces. Considering the net value of interactions for each residue, the Pearson's correlation coefficient between the results of the two methods is 0.634 (p= 6.07E-90) (Fig S2).

Whereas Surfaces does not detect transient interactions, it is possible to generate protein ensembles with the NRGTEN package (Mailhot and Najmanovich, 2021). Such an approach would make possible to generate a distribution of energies for each interaction. Furthermore, the possibility of calculating transition probabilities and occupancies for different configurations opens the possibility to apply statistical mechanics techniques assuming that the ensemble of configurations represents the low energy states that contribute the most to the free energy.

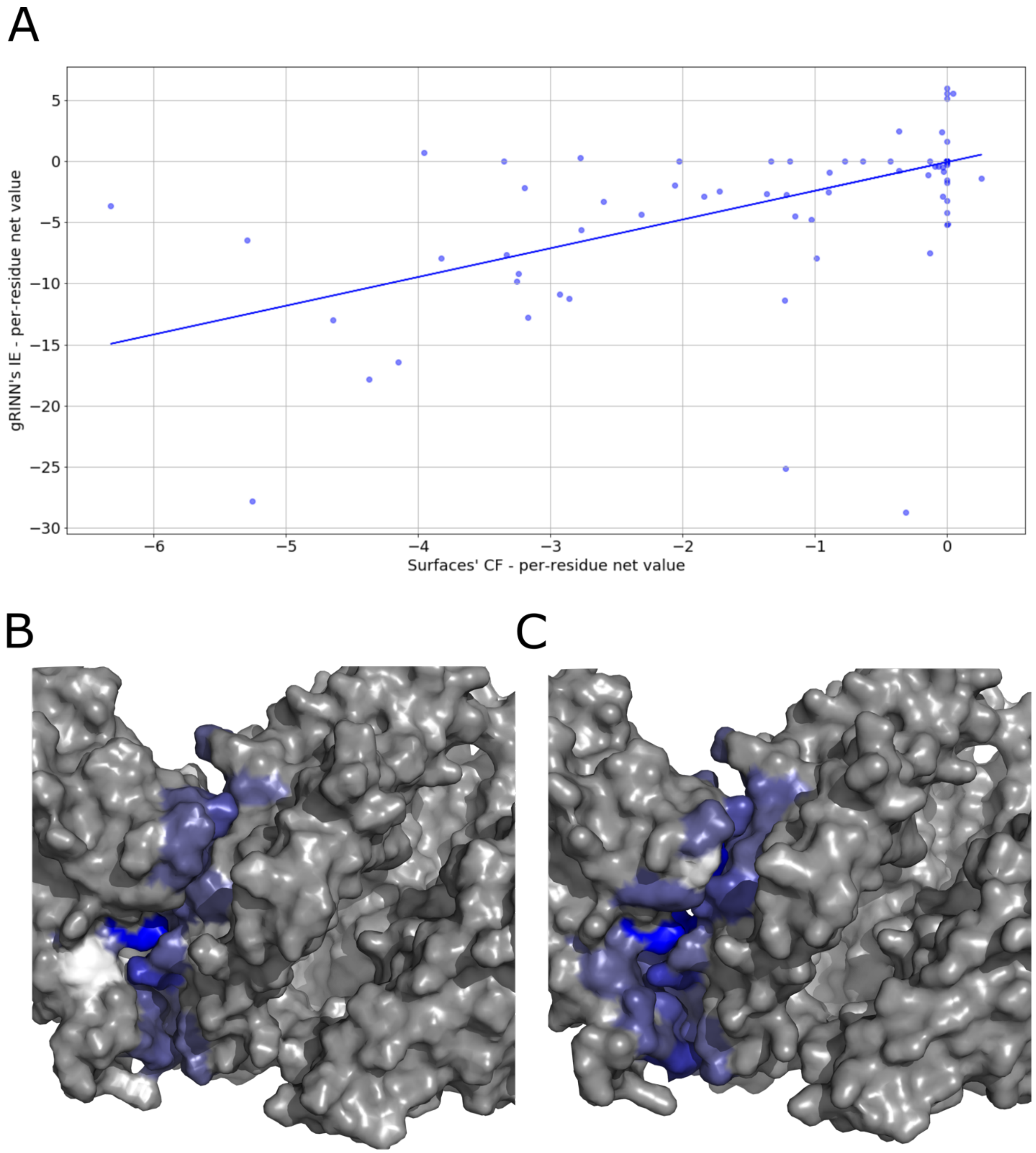


**Fig S2. Per-residue result comparison between gRINN and Surfaces calculations. (A)** Graphic representation of the calculated binding affinities for each residue of the complex using gRINN and Surfaces, as well the tendency line of the correlation between the values. **(B)** Visual representation of the per-residue interactions according to gRINN results in a scale of dark blue (favorable) to white (neutral) generated with Surfaces visual output scripts. **(C)** Visual representation of the per-residue interactions according to Surfaces results in a scale of dark blue (favorable) to white (neutral) generated with Surfaces visual output scripts.

**References**


Barlow,K.A. *et al.* (2018) Flex ddG: Rosetta Ensemble-Based Estimation of Changes in Protein–Protein Binding Affinity upon Mutation. *J Phys Chem B*, 122, 5389–5399.

Bopp,P.A. *et al.* (2008) SCOPE AND LIMITS OF MOLECULAR SIMULATIONS. *Chem Eng Commun*, 195, 1437–1456.

Cameroni,E. *et al.* (2022) Broadly neutralizing antibodies overcome SARS-CoV-2 Omicron antigenic shift. *Nature*, 602, 664–670.

Cheng,M.H. *et al.* (2022) Impact of new variants on SARS-CoV-2 infectivity and neutralization: A molecular assessment of the alterations in the spike-host protein interactions. *iScience*, 25, 103939.

Ciccotti,G. *et al.* (2022) Molecular simulations: past, present, and future (a Topical Issue in EPJB). *Eur. Phys. J. B*, 95, 3.

Dehouck,Y. *et al.* (2013) BeAtMuSiC: prediction of changes in protein–protein binding affinity on mutations. *Nucleic Acids Res*, 41, W333–W339.

Dejnirattisai,W. *et al.* (2022) SARS-CoV-2 Omicron-B.1.1.529 leads to widespread escape from neutralizing antibody responses. *Cell*, 185, 467-484.e15.

Frappier,V. and Najmanovich,R. (2014) Vibrational entropy differences between mesophile and thermophile proteins and their use in protein engineering. *Protein Sci*, 24, 474–483.

Frappier,V. and Najmanovich,R.J. (2014) A Coarse-Grained Elastic Network Atom Contact Model and Its Use in the Simulation of Protein Dynamics and the Prediction of the Effect of Mutations. *PLoS Comput Biol*, 10, e1003569.

Homeyer,N. and Gohlke,H. (2012) Free Energy Calculations by the Molecular Mechanics Poisson−Boltzmann Surface Area Method. *Mol Inform*, 31, 114–122.

Kollman,P.A. *et al.* (2000) Calculating Structures and Free Energies of Complex Molecules: Combining Molecular Mechanics and Continuum Models. *Accounts Chem Res*, 33, 889–897.

Lan,J. *et al.* (2020) Structure of the SARS-CoV-2 spike receptor-binding domain bound to the ACE2 receptor. *Nature*, 581, 215–220.

Mailhot,O. and Najmanovich,R. (2021) The NRGTEN Python package: an extensible toolkit for coarse-grained normal mode analysis of proteins, nucleic acids, small molecules and their complexes. *Bioinformatics*, 37, 3369–3371.

MolSSI COVID-19 Molecular Structure and Therapeutics Hub.

Pahari,S. *et al.* (2020) SAAMBE-3D: Predicting Effect of Mutations on Protein–Protein Interactions. *Int J Mol Sci*, 21, 2563.

Pattnaik,P. (2005) Surface plasmon resonance: applications in understanding receptor-ligand interaction. *Appl Biochem Biotechnol*, 126, 79–92.

Rodrigues,C.H.M. *et al.* (2019) mCSM-PPI2: predicting the effects of mutations on protein–protein interactions. *Nucleic Acids Res*, 47, W338–W344.

Schymkowitz,J. *et al.* (2005) The FoldX web server: an online force field. *Nucleic acids research*, 33, W382-8.

Serçinoglu,O. and Ozbek,P. (2018) gRINN: a tool for calculation of residue interaction energies and protein energy network analysis of molecular dynamics simulations. *Nucleic acids research*, 46, W554–W562.

Sergeeva,A.P. *et al.* (2022) Free energy perturbation calculations of mutation effects on SARS-CoV-2 RBD::ACE2 binding affinity. *Biorxiv*, 2022.08.01.502301.

Teruel,N. *et al.* (2021) Modelling conformational state dynamics and its role on infection for SARS-CoV-2 Spike protein variants. *Plos Comput Biol*, 17, e1009286.

Zhang,N. *et al.* (2020) MutaBind2: Predicting the Impacts of Single and Multiple Mutations on Protein-Protein Interactions. *iScience*, 23, 100939.
